# Mouth-attaching copepod *Salmincola markewitschi* reduces the body condition and growth of juvenile salmonid *Salvelinus leucomaenis* by decreasing host feeding activity: evidence from parasite removal experiment

**DOI:** 10.1101/2024.10.08.617322

**Authors:** Leo Murakami, Ryota Hasegawa, Nozomi Aruga, Nobuhiro Sato, Shingo Nakamura, Hiroshi Kajihara, Itsuro Koizumi

## Abstract

Ectoparasitic copepods in the genus *Salmincola* Wilson, 1915 are considered harmful to host salmonids, yet experimental studies on whether and how these copepods affect hosts are critically limited. We evaluated the effects of the mouth-attaching *Salmincola markewitschi* Shedko and Shedko, 2002 on juvenile white-spotted charr *Salvelinus leucomaenis* (Pallas, 1814) using a parasite removal experiment where all individuals were artificially infected, and then parasites were removed in one group (i.e., treatment), while the copepods were left in another group (i.e., control). We hypothesized that the copepods reduce host feeding, thereby reducing host body condition and growth. The alternative hypothesis is that parasites do not change host feeding, or even increase because of the compensation for the negative effect. In this case, host condition and growth would not be reduced. The control group (parasite remained) had lower food consumption, body condition, and growth compared to the treatment (parasite removed). Thus, *S. markewitschi* negatively affected hosts by reducing host feeding, and no behavioral compensation was observed. It remains unknown, however, whether reduced feeding was due to impaired swallowing or physiological stresses caused by infections. Further experiments on gill- and body-surface-attaching *Salmincola* species could provide deeper insights into how these parasites influence host health.

## Introduction

The genus *Salmincola* Wilson, 1915 comprises a group of ectoparasitic copepods mainly infecting the gills and buccal cavities of freshwater salmonids (Kabata 1969). They are recognized as harmful parasites in aquaculture because their infections cause mechanical damage such as gill lesions and swellings (Kabata and Cousens 1977; Nagasawa et al. 1998; White et al. 2020; Neal et al. 2021). However, quantitative assessments of whether and how these copepods reduce host body condition, growth, and survival are critically lacking, and observational studies in wild populations have shown equivocal results (Chigbu 2001; Nagasawa and Urawa 2002; Ayer et al. 2022; Hasegawa et al. 2022a). Some experimental infections showed a decline in host condition or survival in fish infected with gill-attaching *Salmincola* (Kabata and Cousens 1977; Pawaputanon 1980; Neal et al. 2021), but these did not examine specific mechanisms of health reduction (Pawaputanon 1980; Neal et al. 2021) or lacked proper experimental replication (Kabata and Cousens 1977). In addition, while a negative correlation between host body condition and parasite infection is occasionally reported in the wild (Kusterle et al. 2012; Hasegawa et al. 2022a; Hasegawa and Koizumi 2024), this does not necessarily mean negative effects of parasites, because hosts with poor body condition may be more likely to get infected (e.g., Beldomenico et al. 2008). Overall, the negative effects of *Salmincola* spp. are still largely unknown, and further experimental evidence is required to determine whether and how copepod infections reduce host health.

Here, by conducting a parasite removal experiment, we evaluated whether and how copepods of the genus *Salmincola* affect host salmonids. We hypothesized that *Salmincola* spp. reduce host body condition, growth, and survival by impairing host feeding activities. Parasites generally decrease host activities (Barber et al. 2000) by incurring physiological costs, such as through immune responses (e.g., Sheldon and Verhulst 1996). In addition, mouth-attaching parasites may directly hamper feeding (Vigneshwaran et al. 2019; Hasegawa and Koizumi 2024). The alternative hypothesis is that *Salmincola* spp. would not reduce host health. Host salmonids may compensate for the negative effects of *Salmincola* by increasing foraging and energy intake because host try to recover energy which was declined probably through physiological stress induced by parasites (Hasegawa and Koizumi 2023). We initially infected juvenile (young-of-the-year; hereafter called “YOY”) white-spotted charr *Salvelinus leucomaenis* (Pallas, 1814) with the mouth-attaching *Salmincola markewitschi* Shedko and Shedko, 2002, and the parasites were subsequently removed in one group (i.e., treatment), while they remained in another group (i.e., control). We compared the food consumption, body condition, growth, and survival of white-spotted charr between the two groups. If the parasite negatively affects the host salmonid, body condition, growth, and survival would be reduced. If the negative effect is through hampering of feeding activity, food consumption would decrease. Alternatively, if host salmonid compensate the negative effect of copepods, food consumption would not change or might even increase. In this case, we would predict no significant differences in body condition, growth, and survival between control and treatment groups.

## Methods

*Salmincola markewitschi* naturally occurs in the mouth-cavity of the genus *Salvelinus* Richardson, 1836, particularly white-spotted charr in eastern Russia and Japan (Shedko and Shedko 2002; Nagasawa 2020, 2021; Hasegawa and Koizumi 2024). *Salmincola* spp. have a direct life cycle, which is completed within several months (Kabata and Cousens 1973; Murphy et al. 2020). Free-living copepodids attach to suitable hosts during a short period, generally less than a few weeks (Kabata and Cousens 1973; Murphy et al. 2020). While females embed themselves into the host’s tissue using their “bulla” and generally grow to more than 3–5 mm, males are dwarf forms that attach to the female bodies, and die soon after copulation (Kabata and Cousens 1973; Murphy et al. 2020). Due to this unique life cycle, we focused only females in this study.

Fish were exposed to the parasite at the Sapporo Salmon Museum, Sapporo, Hokkaido, Japan (hereafter called “SSM”), where infections of *S. markewitschi* have been observed since 1985 (Takayama et al. 1999; Nagasawa 2021). In previous studies, we specifically identified these copepods as *S. markewitschi* using both molecular and morphological analyses (Hasegawa et al. 2022b; Shedko et al. 2023), and these specimens and sequences were deposited in the Invertebrate collection of the Hokkaido University Museum (ICHUM 8334) and GenBank (Accession numbers: LC713088, LC713314, and LC713320), respectively. We used YOY white-spotted charr originating from artificial breeding at the SSM. Since these fish were reared in backyards using spring-fed water sources, they had not been exposed to any pathogen infections including *Salmincola* spp. before the experiment.

To expose all experimental fish to infectious copepodids, we placed an enclosure (length × width × height = 1.0 × 1.0 × 0.7 m; mesh size of 20 mm on top and 5 mm on all other sides) in a pond at the SSM. In this pond, many *Salvelinus* spp. were reared and infected with *S. markewitschi* (Nagasawa 2021; Hasegawa et al. 2022b), allowing free-swimming copepodids (less than 1 mm: Murphy et al. 2020) released from these infected fish to freely access and infect the experimental fish in the enclosure.

From 4 August to 15 September 2023, 138 juvenile white-spotted charr were introduced to the enclosure. The enclosure was monitored almost daily, and fish were fed commercial dry pellets (ca. 3.3 mm in diameter, Himesakura 60-3, Higashimaru Co, Ltd). The initial fork length (*L_F_*, to the nearest 1 mm; mean ± standard deviation) and wet body mass (*M*_w_, to the nearest 0.1 g) of introduced fish were 104.7 ± 10.5 mm and 14.3 ± 4.6 g, respectively.

We conducted experimental parasite removal and subsequent fitness evaluations from October to December 2024. These experiments were divided into two trials, which were conducted under the same experimental conditions (Trial 1: 4 October 2023 to 21 November 2023; Trial 2: 16 November to 8 December). At the beginning of each trial, all fish reared in the enclosure were captured and anesthetized by FA100 (Bussan Animal Health Co., Ltd., Osaka, Japan). All fish were briefly examined for copepod infections, including their body surfaces and buccal cavities. In total, 37 infected fish (27 and 10 fish in trial 1 and 2, respectively; see Table 1) were brought to a constant thermostatic chamber and kept individually in tanks (length × width × height; 30 × 20 × 25 cm). The fish were acclimated to laboratory conditions for three days without being fed. The experimental chamber was maintained under constant daylight (12L: 12D) and temperature (14 L) conditions. This temperature range is within the preferred range for *Salmincola* spp. (Yamamoto and Nagasawa 2001; Neal et al. 2021) and white-spotted charr (Takami et al. 1997).

**Table 1.** Summary of biological characters and infection status of the experimental fish.

|  | Treatment (parasite removed) | Control (parasite remained) |
| --- | --- | --- |
| *N.* of fish used | 11 | 11 |
| Initial infection intensity (mean) | 1–2 (1.27) | 1–3 (1.55) |
| Initial $L_F$ (mm) | $125.6 \pm 14.0$ | $124.6 \pm 11.0$ |
| Initial $M_w$ (g) | $23.6 \pm 8.5$ | $21.5 \pm 6.4$ |
| Initial $K$ | $1.15 \pm 0.06$ | $1.08 \pm 0.15$ |
| After $K$ | $1.22 \pm 0.15$ | $1.13 \pm 0.10$ |
| After infection intensity (mean) | NA | 1–3 (1.18) |
| SGR | $0.89 \pm 0.17$ | $0.54 \pm 0.41$ |
| *N.* of dead fish | 0 | 3 |

$L_F$ : Fork length; $M_w$ : Wet body mass; $K$ : Fulton's condition factor; SGR: specific
growth rate. $L_F$ , $M_w$ , $K$ , and SGR are shown with mean ± standard deviation.
Note that two fish individuals in the treatment group died accidentally during the
experiment (they jumped out from the tank).

After acclimation, we recorded the host feeding activity daily for three to five days before the experimental removal. Specifically, we fed each fish commercial dry pellets (see above) once per day and counted the total numbers of pellets consumed as an indicator of host feeding activity (hereafter called “daily consumption”). We dropped five pellets one by one at approximately 3 second intervals in front of the focal fish and waited for about 1 minute. When the focal fish consumed all five pellets, we repeated this process until the fish ceased feeding. If a fish did not consume the first five pellets, the daily consumption was recorded as zero.

After the feeding observations, all fish were measured for *L_F_* and *M*_w_ after being anesthetized with FA100 (after 24-h starvation). We recorded the abundance and attachment locations of all copepods. Then, we randomly assigned the experimental fish to either the control group (without parasite removal) or the treatment group (with parasite removal). For the treatment group, we gently removed all found copepods using forceps. After recovery, we returned all fish to their original tank. There were no significant differences in *L_F_* (Welch’s *t*-test, *t* = −1.717, *d.f.* = 16.17, *p* = 0.105) and *M*_w_ (*t* = 0.436, *d.f.* = 18.00, *p* = 0.668) between the control and treatment groups before the experiment (Table 1). We also found no significant differences in daily consumption between experimental groups (GLMM; *z*-value = 1·416, *p* = 0.157, see below).

After the measurements, each fish was starved for 72 hours, and we resumed recording feeding activities daily as described above. After 32 days, we recorded *L_F_*, *M*_w_, and parasite infections again in the same manner as at the start of the experiment. Throughout the experimental period, we replaced approximately half of the water in the fish tank at least once every four days to maintain clean conditions. We followed all applicable international, national, and institutional guidelines for the care and use of animals.

Of 37 fish (27 and 10 fish in each trial) brought back to the laboratory, copepods died and detached from 5 fish until the start of experiments. Furthermore, 10 fish died (natural mortality: 8 fish, accidental mortality due to jumping out from aquariums: 2 fish) before the first measurements. Therefore, we used 22 fish (control *N* = 11; treatment *N* = 11) for the experiments and subsequent analyses.

Infection indices, such as intensity, were calculated following Bush et al. (1997). To evaluate the impact of the treatments on host feeding activities, we constructed a generalized linear mixed model (GLMM) with a zero-inflated Poisson distribution and a log link function. The response variable was daily consumption and the explanatory variable was the experimental group (i.e., treatment and control). We included fish ID as a random effect.

We compared host body condition (Fulton’s condition factor; *K* = 10^5^ × *M*_w_ / *L* ^3^) and specific growth rate (SGR) between the experimental groups using Welch’s *t*-test. SGR was calculated as follows: (ln *M*_w2_ − ln*M*_w1_)/(d_2_ − d_1_), where *M*_w1_ and *M*_w2_ were the body mass at the start (day _1_) and end (day _2_) of the experiment, respectively. SGRs were also calculated for fish that died during the experiment (*N* = 3), using the date on which death was confirmed as d_2_. Finally, we compared survival between the experimental groups using Fisher’s exact probability test.

All statistical analyses were conducted using R v. 4.0.3 (R Core Team 2021). We performed GLMM with the “*glmmTMB*” package v. 1.1.8 (Brooks et al. 2017). A *p*-value < 0.05 was considered significant, while a *p*-value between 0.05 and 0.10 was considered a meaningful trend. Before conducting *t*-tests, we assessed variance homogeneity with an *F*-test and normality with a Kolmogorov-Smirnov test.

## Results

Of the 22 fish used for the parasite removal experiments (including both control and treatment groups), intensity range was 1–3 (mean 1.41). Half of the fish (64%) had one copepod, while the remaining 36% fish had 2 or 3 copepods. All found copepods were attached to the buccal cavity of the host fish. At the end of the experiment, two fish in the control group (i.e., parasite remained) experienced copepod detachment, but the other 9 fish were still infected with at least one copepod (Table 1).

After the treatment, daily consumption significantly decreased in the control group (i.e., parasites remained) compared to the treatment group (i.e., parasite removed) (GLMM; *z*-value = 2·042, *p* = 0.041; Fig. 1). *K* in the control group was marginally lower compared to the treatment group at the end of the experiment (Welch’s *t*-test; *t* = −1.875, *d.f.* = 19.00, *p* = 0.076, Fig. 1). The SGR of the control group was significantly lower than that of the treatment group (*t* = −2.323, *d.f.* = 13.92, *p* = 0.025, Fig. 1). Three fish died in the control (i.e., parasite remained) group during the experiment (Table 1), whereas no natural mortality was observed in the treatment (i.e., parasite removed) group, except for accidental deaths unrelated to the treatment (two fish died because they jumped out from the tank). There was no significant difference in host survival between experimental groups (Fisher’s exact test; *p* = 0.210).

**Figure 1.**
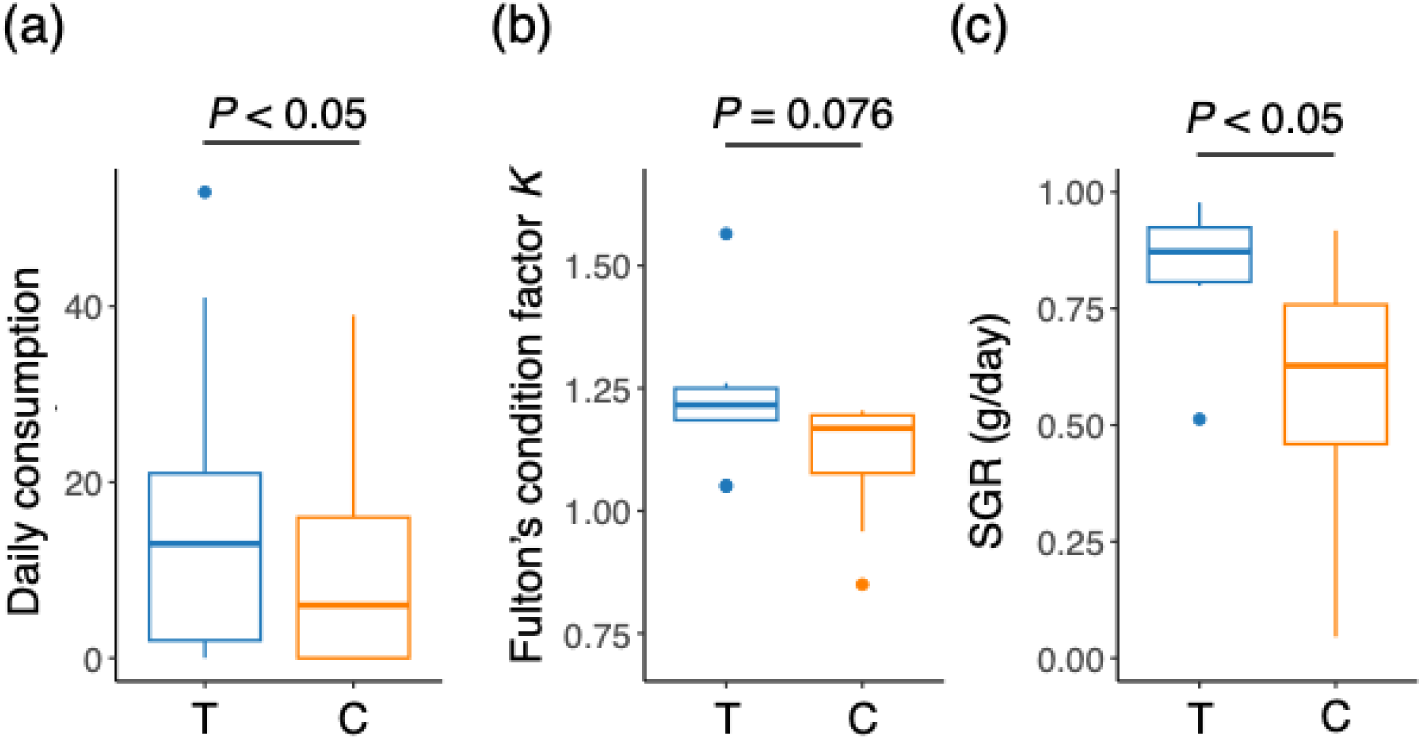
Comparison of (a) daily consumption, (b) Fulton’s condition factor *K*, and (c) specific growth rate (SGR) of juvenile white-spotted charr (*Salvelinus leucomaenis*) between two experimental groups: a treatment group T (in which the copepod *Salmincola markewitschi* was removed) and a control group C (in which the copepod remained).

## Discussion

By conducting the parasite removal experiment, we showed not only the negative effects of *Salmincola* on the host salmonid but also the mechanism by which the parasite reduces host health. Our results supported the first hypothesis that *S. markewitschi* reduces the body condition and growth of juvenile white-spotted charr by reducing host feeding activity. This is consistent with previous observations that *Salmincola* spp. reduce host feeding (Pawaputanon 1980; Nagasawa et al. 1998; Hiramatsu et al. 2001; Roberts et al. 2004); however, experimental studies are particularly important to verify causality. For example, since parasites tend to infect the hosts with poor body conditions (due to a lack of resistance to parasites; Beldomenico et al. 2008), infected hosts observed during a study period might have already been in poor health condition before infection. In this case, parasites are consequences, not causes, of poor body condition, which inevitably leads to a negative correlation between parasite infection and fish body condition (Beldomenico et al. 2008; Beldomenico and Begon 2010). Similarly, a parasite infection experiment without subsequent removal may also introduce a potential bias. That is, experimentally infected fish may be in poorer health or of lower genetic quality, compared to fish that are not infected in the experiments. If this is the case, a simple comparison between infected and uninfected fish cannot account for the innate quality of the individuals. Our removal experiment, on the other hand, excluded these potential biases.

Our removal experiment, however, required relatively complex procedures and time. Therefore, even though we initially introduced more than 100 fish for the artificial infection, our final sample size became relatively small (i.e., 11 treatment and 11 control fish) because many fish remained uninfected, some parasites detached (five out of 37 fish experienced copepod detachment), and some infected fish died (see below). Despite our small final sample size, the effects of *S. markewitschi* were so strong that we detected significant (or marginally significant) decreases in food consumption, body condition, and growth, but not survival. It is likely that the parasite effect on host survival would become significant with a larger sample size and longer experimental periods in the future, considering the negative effects on growth and condition, as well as natural fish mortality occurring only in the control group (*n* = 3, mortality rate 27%). It is worth mentioning that the fish that showed natural mortality before the experiment were all infected with copepods (eight out of 37 fish brought back to the laboratory; mortality rate 22%) and these fish showed a decline in feeding activity (Murakami, personal observations), suggesting that *Salmincola* infections have deleterious impacts on survival through loss of feeding. In fact, a previous infection experiment showed that gill-attaching *S. californiensis* (Dana, 1852) caused high mortality (48%) in juvenile kokanee salmon *Oncorhynchus nerka* (Walbaum, 1792) (Kabata and Cousens 1977).

Our alternative hypothesis (i.e., compensatory foraging) was rejected. We previously showed through field observation that some white-spotted charr infected with *S. markewitschi* were more likely to be caught by angling, suggesting that they compensate for parasite infection by increasing energy intake (Hasegawa and Koizumi 2023). This may because host holding resources (energy) was declined through physiological stress induced by parasites (Hasegawa and Koizumi 2023). The discrepancy can be explained by differences in fish size and condition. The infected fish caught by angling tended to be larger and in better body condition because they likely had enough energy to compensate (Hasegawa and Koizumi 2023). The fish used in the current experiment were all small (i.e., YOY), and thus, no extra-energy would have remained.

Mouth-attaching *Salmincola* should have a greater impact compared to gill-attaching or body-surface-attaching *Salmincola* because they impair host feeding both mechanically and physiologically. Indeed, we occasionally observed that some infected fish could not swallow pellets (Murakami, unpublished data), possibly due to blockage caused by the parasite or diminished suction power resulting from reduced oral cavity space. In addition, infections of *Salmincola* cause physical damage at attachment sites (Kabata and Cousens 1977; Nagasawa et al. 1998; White et al. 2020), and host may try to minimize their energy expenditures and reduce feeding activities to recover this damage (Brassard et al. 1982; Øverli et al. 2014). Host fish may also enhance their immunity to fight against parasite infections, which may also reduce overall host activities (Øverli et al. 2014; Jolles et al. 2020). Understanding the relative contributions of mechanical and physiological stress is essential for predicting the broader impacts of the genus *Salmincola*. Reduced host feeding activity has also been reported in some gill-attaching *Salmincola* (Pawaputanon 1980; Roberts et al. 2004), indicating that physiological stress alone negatively affects host health. In addition, gill-attaching *Salmincola* alter host behavior in ways that facilitate further infections (Poulin et al. 1991). Further experimental studies are needed to reveal the detailed mechanisms by which *Salmincola* affects salmonid hosts.

Significant declines in feeding, body condition, and growth rate caused by *Salmincola* could have substantial effects on host demographic features, such as host abundance and dynamics. Indeed, epizootics of *Salmincola* have recently become major problems in North America, and several studies have raised concerns about host population-level extirpations caused by *Salmincola* (Monzyk et al. 2015; Mitro 2016; Lepak et al. 2022). Most previous studies provided ambiguous evidence of the negative effects of the copepods on host fitness traits in the wild (Chigbu, 2001; Nagasawa and Urawa 2002; Ayer et al., 2022), and some researchers have concluded that their negative effects are negligible in wild populations because infection levels are generally low (Black et al. 1983; Amundsen et al. 1997). In contrast, our study provided firm evidence of a loss in host condition, growth, and potentially survival, even when fish were infected with only a few copepods, which is similar to infection levels in wild populations (Hasegawa and Koizumi 2021, 2024), indicating that their impact on host health in the wild may be stronger than previously considered. Therefore, continuous census and monitoring are necessary to evaluate population-level consequences of salmonid-*Salmincola* systems, which is also useful for host conservation and aquaculture.

## Acknowledgements

Dr. Koh Hasegawa kindly provided us an enclosure. Staff and volunteers of Sapporo Salmon Museum provided us experimental fish and technically supported our experiments. Natsuki Hara and Hironori Mieda helped our experiments. Drs Keiichi Kakui and Robert Poulin provided critical comments on our earlier drafts. This study was partially supported by the Japan Society for the Promotion of Science Research Fellow Grant (Grant No. JP2J12519 to RH).

## Data availability statement

Data can be provided by the corresponding author upon reasonable request.

## Authors contributions

Conceptualization: LM, RH, NA, NS, SN, HK, IK

Data curation: LM, RH

Formal analysis: LM, RH

Funding acquisition: RH

Investigation: LM, RH, NA, NS, SN, IK

Supervision: RH, HK, IK

Visualization: RH

Writing-original draft: LM, RH

Writing-review & editing: LM, RH, NA, NS, SN, HK, IK

